# Interaction of a sarcolipin pentamer and monomer with the sarcoplasmic reticulum calcium pump, SERCA

**DOI:** 10.1101/696070

**Authors:** J. P. Glaves, J. O. Primeau, P. A. Gorski, L. M. Espinoza-Fonseca, M. J. Lemieux, H. S. Young

## Abstract

The sequential rise and fall of cytosolic calcium underlies the contraction-relaxation cycle of muscle cells. While contraction is initiated by the release of calcium from the sarcoplasmic reticulum, muscle relaxation involves the active transport of calcium back into the sarcoplasmic reticulum. This re-uptake of calcium is catalysed by the sarco-endoplasmic reticulum Ca^2+^-ATPase (SERCA), which plays a lead role in muscle contractility. The activity of SERCA is regulated by small membrane protein subunits, most well-known being phospholamban (PLN) and sarcolipin (SLN). SLN physically interacts with SERCA and differentially regulates contractility in skeletal and atrial muscle. SLN has also been implicated in skeletal muscle thermogenesis. Despite these important roles, the structural mechanisms by which SLN modulates SERCA-dependent contractility and thermogenesis remain unclear. Here, we functionally characterized wild-type SLN and a pair of mutants, Asn^4^-Ala and Thr^5^-Ala, which yielded gain-of-function behavior comparable to what has been found for PLN. Next, we analyzed twodimensional crystals of SERCA in the presence of wild-type SLN by electron cryo-microscopy. The fundamental units of the crystals are anti-parallel dimer ribbons of SERCA, known for decades as an assembly of calcium-free SERCA molecules induced by the addition of decavanadate. A projection map of the SERCA-SLN complex was determined to a resolution of 8.5 Å, which allowed the direct visualization of a SLN pentamer. The SLN pentamer was found to interact with transmembrane segment M3 of SERCA, though the interaction appeared to be indirect and mediated by an additional density consistent with a SLN monomer. This SERCA-SLN complex correlated with the ability of SLN to decrease the maximal activity of SERCA, which is distinct from the ability of PLN to increase the maximal activity of SLN. Protein-protein docking and molecular dynamics simulations provided models for the SLN pentamer and the novel interaction between SERCA and a SLN monomer.

**STATEMENT OF SIGNIFICANCE:** This research article describes a novel complex of the sarcoplasmic reticulum calcium pump SERCA and its regulatory subunit sarcolipin. Given the potential role of sarcolipin in skeletal muscle non-shivering thermogenesis, the interactions between SERCA and sarcolipin are of critical importance. Using complementary approaches of functional analysis, electron crystallography, and molecular dynamics simulations, we demonstrate an inherent interaction between SERCA, a sarcolipin monomer, and a sarcolipin pentamer. The interaction involves transmembrane segment M3 of SERCA, which allows sarcolipin to decrease the maximal activity or turnover rate of SERCA. Protein-protein docking and molecular dynamics simulations provided models for the SLN pentamer and the novel interaction between SERCA and a SLN monomer.

## INTRODUCTION

A major regulator of cellular calcium homeostasis in skeletal muscle is the sarco-endoplasmic reticulum Ca^2+^-ATPase (SERCA). Calcium release channels trigger muscle contraction by releasing calcium stored in the sarcoplasmic reticulum (SR). In turn, the SERCA pump (SERCA1a isoform) triggers muscle relaxation by returning calcium from the cytosol to the lumen of the SR. In skeletal and atrial muscle, SERCA is regulated by sarcolipin (SLN), which allows for dynamic control of calcium homeostasis and the contraction-relaxation cycle. SLN is a 31-residue tail-anchored integral membrane protein that resides in the SR membrane and acts as an inhibitor of SERCA. SLN is homologous to phospholamban (PLN), a well-known regulatory subunit of SERCA in cardiac muscle (SERCA2a isoform). Both regulatory subunits inhibit SERCA by lowering its apparent affinity for calcium. The relief of inhibition and re-activation of SERCA is mediated by adrenergic signaling pathways and the reversible phosphorylation of PLN and SLN, which in turn has significant effects on calcium uptake in muscle tissues.

Over the last few years, studies have revealed that SLN belongs to a family of regulatory subunits that target SERCA in a tissue-specific manner, herein collectively described as the “regulins” (1–3). Prior to this, PLN and SLN were the only known SERCA regulatory subunits, with the importance of PLN in cardiac muscle overshadowing the role of SLN in skeletal muscle. However, SLN is also found in atrial muscle, alongside PLN, where it plays an important, but undefined role in cardiac contractility. In addition, interest in SLN has intensified with the identification of a new physiological role – SLN appears to be involved in non-shivering, skeletal muscle-based thermogenesis (4). The current theory is that SLN promotes uncoupling of SERCA, where the net balance of ATP hydrolysis does not correlate with productive calcium transport. The excess ATP hydrolysis contributes to thermal energy in skeletal muscle, the largest tissue mass in the human body. Given the abundance of skeletal muscle, and the role of SLN in thermogenesis and energy balance, SLN may also contribute to metabolic disorders such as obesity and diabetes. With this potential new functionality of SLN, a molecular understanding of the SERCA regulatory mechanism is imperative.

The homology between SLN and PLN lies within their transmembrane domains, and this has long suggested commonality in function. However, the structure of SLN is quite distinct, with a short cytoplasmic domain (residues 1-7), a transmembrane α-helix (residues 8-26), and a unique luminal tail (residues 27-31). Like PLN, SLN alters the apparent calcium affinity of SERCA, yet there are substantial differences in the mechanism (5). The inhibitory properties of SLN are strongly dependent on the highly-conserved C-terminal tail (Arg^27^-Ser-Tyr-Gln-Tyr^31^ or RSYQY sequence), whereas the inhibitory properties of PLN are encoded in its transmembrane domain. In addition, SLN appears to remain associated with SERCA throughout the calcium transport cycle (6). Recall that PLN is thought to interact with calcium-free forms of SERCA and dissociate from the enzyme under certain conditions (e.g. elevated calcium plus PLN phosphorylation). Crystal structures of a SERCA-SLN complex have been determined, revealing SLN binding to the inhibitory groove (M2, M6, & M9) in a novel E1-like state of SERCA (7,8). This novel conformation of SERCA was not the anticipated calcium-free E2 state and the luminal RSYQY sequence of SLN was not in direct contact with SERCA. Thus, it seems likely that the crystal structures represent one intermediate as SERCA and SLN progress together through the calcium transport cycle, and that the SERCA-SLN complex involves multiple conformational states. How the regulatory RSYQY sequence of SLN interacts with SERCA remains elusive.

We have previously shown that PLN pentamers interact with SERCA in the membrane environment of large two-dimensional (2D) co-crystals, and that the association is distinct from the inhibitory interaction and dependent on the functional state of PLN (9,10). Our recent study compared helical crystals of the SERCA-PLN complex with the large 2D crystals, which allowed us to conclude that the PLN pentamer naturally associates with SERCA at a distinct site (11). This interaction correlated with the ability of PLN pentamers to increase the maximal activity of SERCA at high calcium concentrations, and a molecular model of the novel SERCA-PLN complex was presented. Given the similar functional properties of SLN and observations that SLN can form oligomers (12,13), we set out to determine if SLN could be co-crystallized with SERCA. Here, we functionally characterized wild-type SLN, as well as gain of function mutants Asn^4^-Ala and Thr^5^-Ala (**Figure 1**). These mutants confirm structural and functional elements that are conserved in PLN and SLN. In addition, we analyzed large 2D crystals of the SERCA-SLN complex by electron cryo-microscopy. Both wild-type SLN and a gain-of-function mutant (Asn^4^-Ala) were evaluated. We report the first direct observation of a SLN pentamer and we show that it interacts with SERCA in a manner analogous to that previously observed for PLN. However, instead of a direct interaction between the SLN pentamer and transmembrane segment M3 of SERCA, this interaction is mediated by an additional density most consistent with a SLN monomer. The ability of SLN and PLN to interact with an accessory site of SERCA (M3) appears to contribute to the regulatory mechanism. However, the interactions of the PLN and SLN pentamers with SERCA are different and the functional effects on the maximal activity of SERCA are opposite – PLN increases while SLN decreases the maximal activity of SERCA.

**Figure 1:**
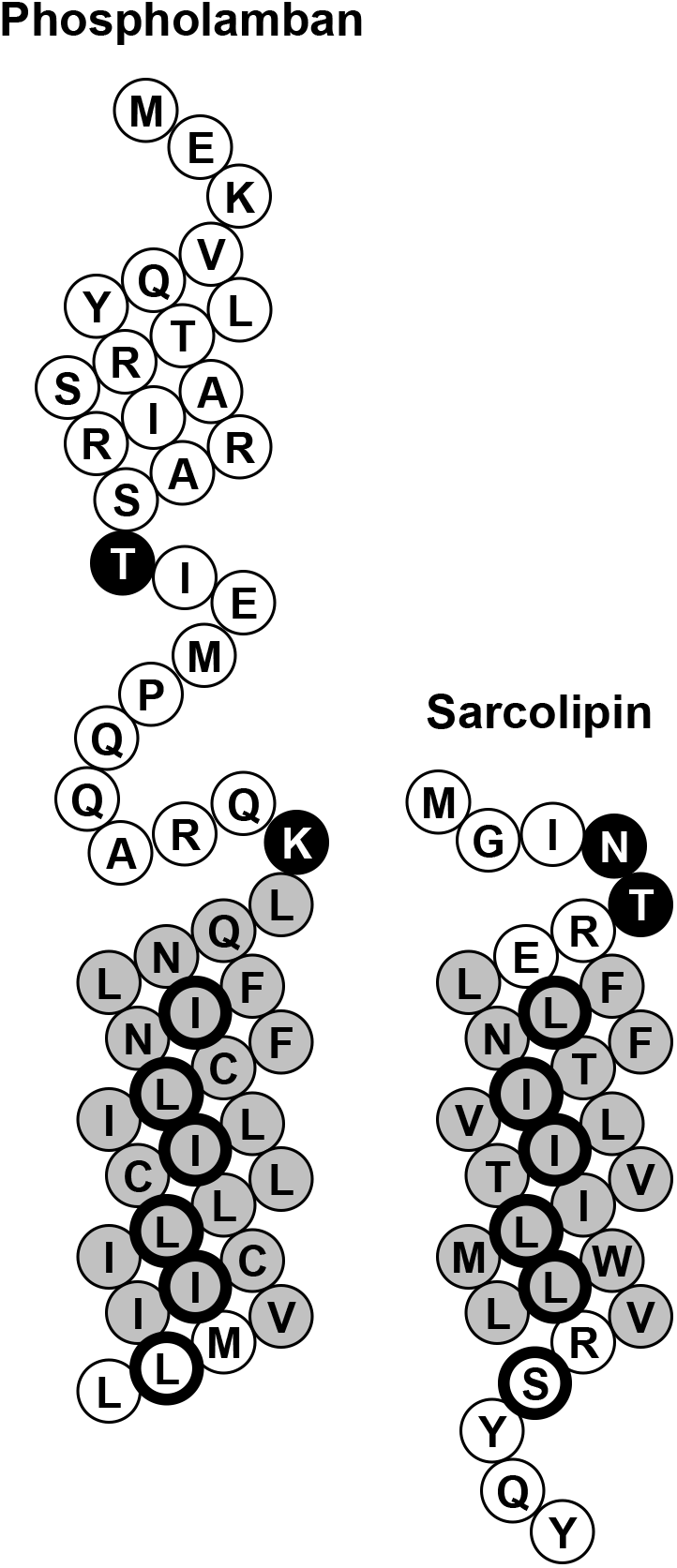
Topology diagrams for human phospholamban (PLN) and sarcolipin (SLN). The transmembrane domains are colored grey and the cytoplasmic and luminal domains are colored white. Asn^4^ and Thr^5^ are indicated in black, as are the comparative residues in PLN (Lys^27^ and Thr^17^). The Leu-Ile residues of PLN that stabilize the pentameric state are outlined in black (Ile^33^, Leu^37^, Ile^40^, Leu^44^, Ile^47^, and Leu^51^), as are the comparative residues in SLN (Leu^10^, Ile^14^, Ile^17^, Leu^21^, Leu^24^, and Ser^28^).

## MATERIALS & METHODS

### Materials

The following reagents were of the highest purity available: octaethylene glycol monododecyl ether (C_12_E_8_; Barnet Products, Englewood Cliff, NJ); egg yolk phosphatidylcholine (EYPC), phosphatidylethanolamine (EYPE) and phosphatidic acid (EYPA) (Avanti Polar Lipids, Alabaster, AL); reagents used in the coupled enzyme assay (Sigma-Aldrich, Oakville, ON Canada).

### Expression and purification of recombinant SLN

Recombinant SLN was expressed and purified as previously described (14) with an additional organic extraction step. Following protease digestion of the maltose-binding protein and SLN fusion protein, trichloroacetic acid was added to a final concentration of 6%. This mixture was incubated on ice for 20 minutes. The precipitate was collected by centrifugation at 4°C and subsequently homogenized in a mixture of chloroform-isopropanol-water (4:4:1) and incubated at room temperature for 3 hours. The organic phase, highly enriched in recombinant SLN, was removed, dried to a thin film under nitrogen gas and resuspended in 7 M GdnHCl. Reverse-phase HPLC was performed as described (14).

### Co-reconstitution of SLN and SERCA

Lyophilized SLN (30 μg or 75 μg) was resuspended in a 75 μl mixture of chloroform-trifluroethanol (2:1) and mixed with lipids (400 μg EYPC, 50 μg EYPE, 50 μg EYPA) from stock chloroform solutions. The peptide-lipid mixture was dried to a thin film under nitrogen gas and desiccated under vacuum overnight. The peptide-lipid mixture was hydrated in buffer (20 mM imidazole pH 7.0; 100 mM KCl; 0.02% NaN_3_) at 37 °C for 10 min, cooled to room temperature, and detergent-solubilised by the addition of C_12_E_8_ (0.2 % final concentration) with vigorous vortexing. Detergent-solubilized SERCA was added (500 μg in a total volume of 200 μl) and the reconstitution was stirred gently at room temperature. Detergent was slowly removed by the addition of SM-2 biobeads (Bio-Rad, Hercules, CA) over a 4-hour time course (final ratio of 25 biobeads: 1 detergent w/w). Following detergent removal, the reconstitution was centrifuged over a 20-50% sucrose step-gradient for 1 h at 100,000g. The resultant layer of reconstituted proteoliposomes was removed, flash-frozen in liquid-nitrogen and stored at −80 °C. The final approximate molar ratios were 120 lipid : 2 or 5 SLN : 1 SERCA (15).

### ATPase activity assays of SERCA reconstitutions

ATPase activity of the co-reconstituted proteoliposomes was measured by a coupled-enzyme assay over a range of calcium concentrations from 0.1 μM to 10 μM (15,16). The Kca (apparent calcium affinity) and V_max_ (maximal activity) were determined by fitting the data to the Hill equation (Sigma Plot software, SPSS Inc., Chicago, IL). Errors were calculated as the standard error of the mean for a minimum of three independent reconstitutions.

### Crystallization of SLN with SERCA

SLN and SERCA were co-reconstituted in the presence of a lipid mixture (final lipid ratio of 8 EYPC : 1 EYPA : 1 EYPE) that promoted vesicle fusion and crystallization (10,17,18). Co-reconstituted proteoliposomes were collected by centrifugation in crystallization buffer (20 mM imidazole pH 7.0, 100 mM KCl, 35 mM MgCl2, 0.5 mM EGTA, 0.25 mM Na3VO4, 30 μM thapsigargin). Note that the sodium orthovandate (Na3VO4) was converted to decavanadate and immediately added to crystallization buffer (11). The samples were subjected to four freeze-thaw cycles, followed by incubation at 4°C for up to one week. Three to five days were optimal for the highest frequency and quality of two-dimensional crystals. For initial screening of crystals by negative stain, crystals were adsorbed to carbon-coated grids and stained with 2% uranyl acetate. The grids were blotted with filter paper and air-dried. Crystals suitable for imaging were adsorbed to holey-carbon grids and plunge-frozen in liquid ethane.

### Electron microscopy of frozen hydrated SERCA and SLN co-crystals

Crystals were imaged in a Tecnai F20 electron microscope (FEI Company, Einhoven, Netherlands) in the Microscopy and Imaging Facility (University of Calgary) or a JEOL 2200FS electron microscope (JEOL Ltd., Tokyo, Japan) in the Electron Microscopy Facility (National Institute for Nanotechnology, University of Alberta and National Research Council of Canada). A standard room temperature holder was used for negatively stained samples and a Gatan 626 cryoholder (Gatan Inc., Pleasanton, CA) was used for frozen-hydrated samples. The microscope was operated at 200 kV and low-dose images were recorded at a magnification of 50,000x. The best films were digitized at 6.35 μm/pixel with a Nikon Super Coolscan 9000 followed by pixel averaging to achieve a final resolution of 2.54 Å/pixel. All data were recorded with defocus levels of 0.5-2 μm with an emphasis on low-defocus images for frozen-hydrated samples (0.5 to 1.0 μm).

### Data processing

The MRC image processing suite was used for images of frozen-hydrated SERCA-SLN crystals (19). Two rounds of unbending were performed prior to extracting amplitudes and phases from each image. Data was then corrected for the contrast transfer function using the program PLTCTFX (20). Common phase origins for merging were determined in the p22_1_2_1_ plane group using ORIGTILT with reflections of signal-to-noise ratio (IQ) <4. For averaging, data were weighted based on IQ including data with IQ <7, and the corresponding phase residuals represent the inverse cosine of the figure of merit from this averaging. Projection maps were calculated by Fourier synthesis from the averaged data using the CCP4 software suite (21).

### Molecular modeling of SLN pentamer and SLN bound to SERCA

The hybrid solution and solid-state NMR structure of the PLN pentamer (PDB accession code 2KYV) and the structure of SLN (PDB accession code 3W5A) were used as templates for the construction of an atomistic model of the SLN pentamer. SLN monomers were aligned to each PLN monomer position in the structure of the PLN pentamer. The resultant model of the SLN pentamer was subjected to 5,000 steps of energy minimization.

The crystal structure of SERCA1a in the E2·MgF_4_^2-^ state (PDB accession code 1WPG) and the structure of SLN (PDB accession code 3W5A) were used as templates for the construction of an atomistic model of the SERCA-SLN complex. The model of the SERCA-SLN complex was guided by the relative positions of SERCA and the SLN monomer in projection maps of the large 2D crystals (**Figure 5**). Protein-protein docking simulations identified an appropriate binding interface between the M3 helix of SERCA and the SLN monomer using ClusPro (22). The resulting docked orientations were clustered and compared against the 2D crystallographic data to select the most appropriate M3-SLN complex. The model of the SERCA-SLN complex was then subjected to 5,000 steps of energy minimization.

### Molecular dynamics simulations of the SLN pentamer and SERCA-SLN complex

The SLN pentamer and SERCA-SLN complex served as starting models to obtain structures of these complexes at physiologically relevant simulation conditions. Based on previous studies of the E2 state of SERCA (23), we modeled transport site residues Glu^309^, Glu^771^ and Glu^908^ as protonated and residue Asp^800^ as ionized. Both complexes were inserted in a pre-equilibrated 150 Å × 150 Å bilayer of POPC:POPE:POPA lipids (8:1:1). This lipid-protein system was solvated using TIP3P water molecules with a minimum margin of 15 Å between the protein and the edges of the periodic box in the z-axis, and K+ and Cl^−^ ions were added (KCl concentration of ~0.1 mM). Molecular dynamics (MD) simulations of the fully solvated complexes were performed using NAMD 2.12 (24) with periodic boundary conditions (25), particle mesh Ewald (26,27), a nonbonded cut-off of 9 Å, and a 2 fs time step. The CHARMM36 force field topologies and parameters were used for the proteins (28), lipids (29), waters, and ions. The NPT ensemble was maintained with a Langevin thermostat (310K) and an anisotropic Langevin piston barostat (1 atm). The systems were first subjected to energy minimization, followed by gradually warming up of the system for 200 ps. This procedure was followed by 10 ns of equilibration with backbone atoms harmonically restrained using a force constant of 10 kcal mol^−1^ Å^−2^. The MD simulations of the complexes were continued without restraints for 400 ns (**Supplemental Figure 1**).

## RESULTS

Since SLN is a homolog of PLN found in skeletal muscle and the atria of the heart, it was reasonable to consider whether SLN can interact with transmembrane segment M3 of SERCA. While SLN and PLN have a similar inhibitory effect on SERCA, the molecular mechanisms by which they regulate SERCA are distinct. The inhibitory properties of PLN are encoded in the transmembrane domain, whereas the inhibitory properties of SLN are strongly influenced by its unique luminal domain (5). The cytoplasmic domains of both PLN and SLN allow for reversal of SERCA inhibition by phosphorylation, though the signaling pathways and molecular mechanisms are different. In addition, PLN is known to form a pentamer that persists in SDS-PAGE analyses, whereas SLN has a reduced tendency to form higher-order oligomers with monomers and dimers being the prevalent species identified by SDS-PAGE (12). Thus, the interaction of an oligomeric species (e.g. pentamer) with SERCA may not occur or may be different for SLN. To investigate this, we co-reconstituted SERCA with wild-type and mutant forms of SLN. ATPase activity of the co-reconstituted proteoliposomes demonstrated a functional interaction between the proteins, and we correlated these data with structural analyses from 2D crystals and molecular dynamics (MD) simulations.

### Co-reconstitution of SERCA and SLN

In the present study, we produced proteoliposomes containing a high density of SERCA and SLN with a lipid-to-protein molar ratio of approximately 120-to-1 and a SERCA-SLN molar ratio of either 1:2 or 1:5 (9,14,15,30,31). The 1:5 SERCA-SLN ratio was used throughout, with the exception of the 1:2 ratio used for comparative activity assays (see below). Focusing on wild-type and two mutant forms of SLN, we measured the calcium-dependent ATPase activity of SERCA in the absence and presence of SLN. The measurement of ATP hydrolysis rates by reconstituted SERCA proteoliposomes yielded an apparent calcium affinity (K_Ca_) of 0.41 ± 0.02 μM for SERCA alone and 0.76 ± 0.02 μM for SERCA in the presence of wild-type SLN (**Figure 2A**). This level of inhibitory activity is consistent with earlier SLN and SERCA co-reconstitution (12,14,32) and heterologous co-expression studies (33). We also studied two mutants of SLN, a previously characterized gain-of-function mutant (Thr^5^-Ala; not shown) and an uncharacterized mutant (Asn^4^-Ala; **Figure 2A**) designed to mimic the Lys^27^-Ala gain-of-function mutant at the homologous position in human PLN (9). Asn^4^-Ala and Thr^5^-Ala were both gain-of-function mutants, further reducing the apparent calcium affinity of SERCA (K_Ca_ values were 1.27 ± 0.05 and 0.95 ± 0.05 μM, respectively). Thus, the potent gain of function observed for the Asn^4^-Ala mutant supported the notion that Asn^4^ of human SLN and Lys^27^ of human PLN (**Figure 1**) serve the analogous function of influencing the stability of the inhibitory complex (34). In addition, the gain of function observed for the Thr^5^-Ala mutant compares well with a similar mutation in PLN, Thr^17^-Ala, which has been reported to be a gain-of-function form of PLN (35).

**Figure 2:**
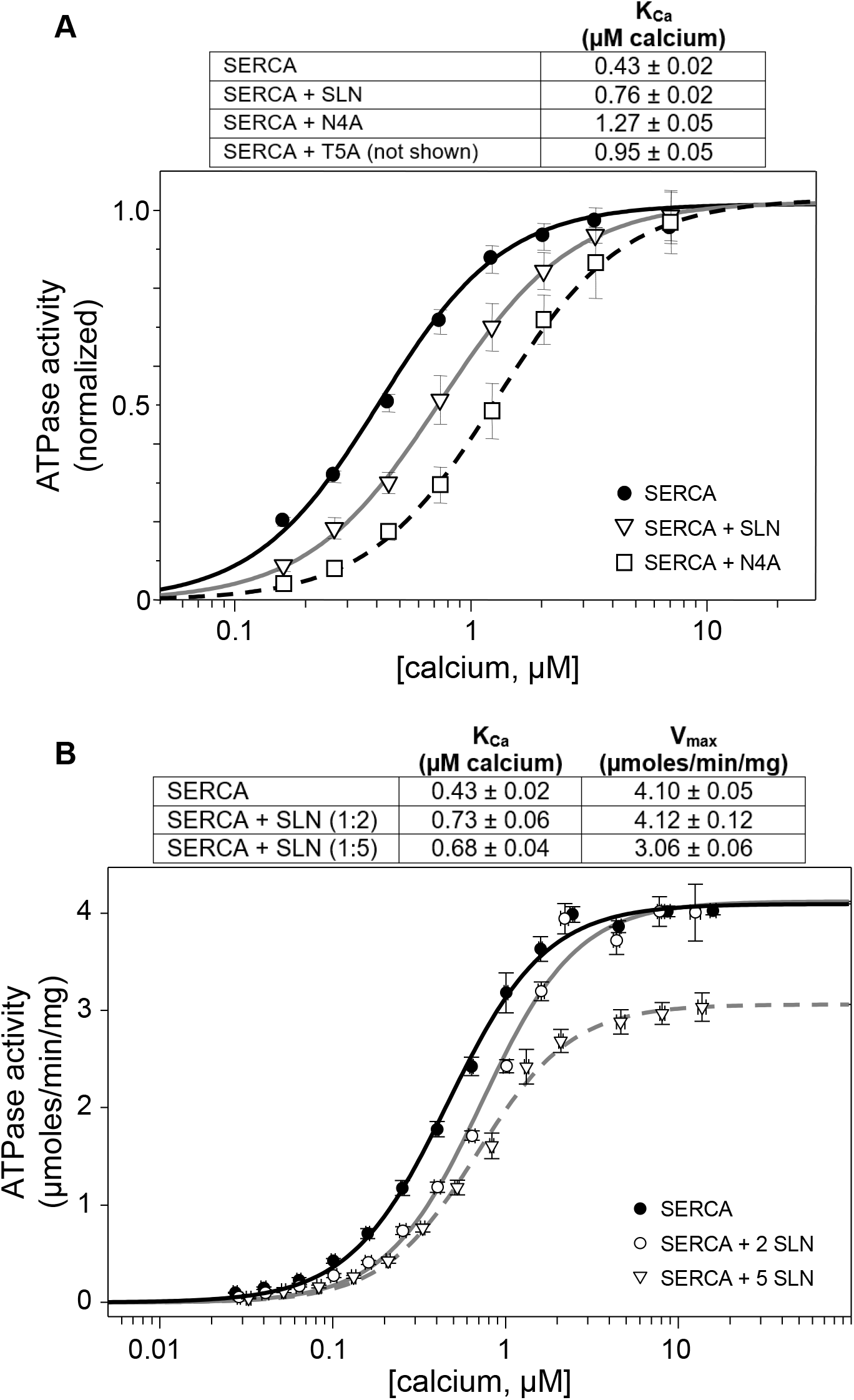
ATPase activity of co-reconstituted proteoliposomes containing SERCA and SLN. **(A)** ATPase activity of SERCA reconstituted in the absence (SERCA; *circles*) and in the presence of wild type SLN (SERCA + SLN; *triangles*) and Asn^4^-Ala SLN (SERCA + N4A; *squares*) at a 1:5 molar ratio. The calcium affinity values (K_Ca_) are shown in the inset table, and all curves have been normalized to the maximal activity (V/V_max_). Notice the gain-of-function behaviour observed for the Asn^4^-Ala mutant (larger rightward shift of the ATPase activity curve). **(B)** ATPase activity of SERCA reconstituted in the absence of SLN (*black circles*), in the presence of 1:2 molar ratio (*white circles*) and a 1:5 molar ratio (*triangles*) of SERCA to wild-type SLN. The calcium affinity (K_Ca_) and maximal activity (V_max_) values are shown in the inset table. Notice the decrease in the V_max_ of SERCA at the 1:5 molar ratio of SERCA-SLN.

The effect of PLN on the apparent calcium affinity of SERCA saturates at a molar ratio of approximately one PLN monomer per SERCA (36,37) and it is assumed that SLN behaves in a similar manner. However, the PLN pentamer also affects the maximal activity of SERCA in a concentration dependent manner (11). This raises the question what are the consequences of higher ratios of SLN on SERCA maximal activity? To address this, we measured the calcium-dependent ATPase activity of SERCA alone and in the presence of two molar ratios of SERCA-SLN (**Figure 2B**). The SERCA-SLN ratios were designed to saturate an effect on the apparent calcium affinity of SERCA (1:2 ratio), and uncover any effects on the maximal activity of SERCA (1:5 ratio). Similar to what was observed for PLN, the effect of SLN on the apparent calcium affinity (K_Ca_) of SERCA remained unchanged for the two molar ratios. In contrast to what was observed for PLN, there was a statistically significant decrease in the maximal activity (V_max_) of SERCA at the 1:5 molar ratio of SERCA-SLN. The V_max_ value for SERCA alone (4.10 ± 0.05 μmoles/min/mg) was similar to SERCA in the presence of the 1:2 ratio of SLN (4.12 ± 0.12 μmoles/min/mg). In contrast, the decreased V_max_ for SERCA in the presence of the 1:5 ratio of SLN (3.06 ± 0.06 μmoles/min/mg) was statistically significant from SERCA alone and SERCA in the presence of the 1:2 ratio of SLN (p<0.01). This result supports the notion that SERCA activity is influenced by the membrane concentration of SLN, though the effect is opposite to that seen for PLN. The decrease of SERCA’s V_max_ occurs at the higher concentration of SLN in the membrane, suggesting that it may involve an oligomeric form of SLN and a distinct interaction between SLN and SERCA.

### Two-dimensional co-crystals of SERCA and SLN

Proteoliposomes containing SERCA in the presence of a 1:5 molar ratio of wild-type and mutants forms of SLN were capable of forming large 2D crystals (9,30). Interestingly, wild-type SLN generally produced fewer co-crystals with SERCA than the Asn^4^-Ala gain-of-function mutant of SLN. A similar trend was observed for wild-type PLN and the Lys^27^-Ala gain-of-function mutant (9). Despite the differences in crystal frequency, the morphology and lattice parameters were comparable to one another and to those previously reported for SERCA-PLN crystals (p22_1_2_1_ plane group symmetry; *a* ~ 346 Å and *b* ~ 70 Å (9,30)). Given that the fundamental units of the crystals are SERCA dimer ribbons rigidly held together by decavanadate (38,39), this was an indication that PLN and SLN may have a similar mode of interaction with SERCA. For wild-type SLN, a projection map of negatively stained 2D crystals was calculated after averaging Fourier data from at least five independent crystals (~20 Å resolution). We used standard methods for correcting lattice distortions, and no CTF correction was applied. The projection map from negatively stained crystals revealed SERCA dimer ribbons as a fundamental feature of the crystals, with densities consistent with SLN oligomers interspersed between the SERCA dimer ribbons (**Figure 3**). The relative size and position of the SLN oligomer density was similar to PLN. We concluded that an oligomeric form of SLN could interact with SERCA in the 2D co-crystals, though the negative stain projection maps did not reveal the size and stoichiometry of the complex.

**Figure 3:**
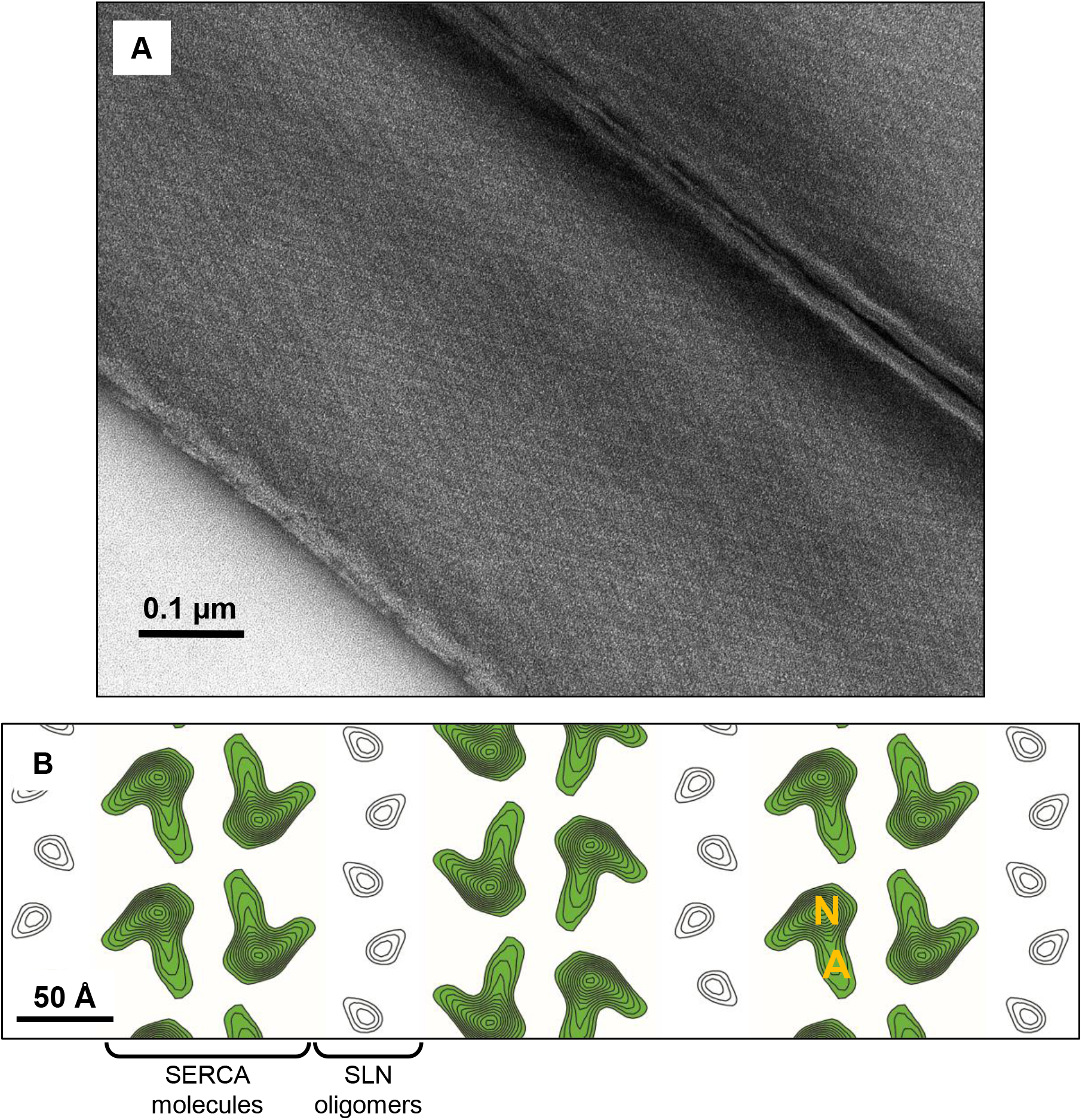
Two-dimensional crystals of SERCA and wild-type SLN. **(A)** Negatively stained image of two side-by-side crystals. The central crystal is a flattened cylinder approximately 0.5 μm wide, with the two independent lattices (one from each side of the crystal) appearing as diagonal arrays in the image. The scale bar is 0.1 μm. **(B)** Projection map from negatively stained 2D crystals of SERCA and wild-type SLN (SERCA density is highlighted in *green*; SLN density is *white* contours). The projection map is contoured showing only positive (continuous lines) densities; each contour level corresponds to 0.25 σ. The relative positions of the nucleotide binding (N) and actuator (A) domains of SERCA are indicated.

To achieve this level of detail, frozen-hydrated specimens were used for the structural characterization of a SERCA-SLN complex in a membrane environment. In the case of PLN (9), we focused on co-crystals of the Lys^27^-Ala gain-of-function mutant because the abundance of crystals facilitated data collection. Indeed, we initially pursued the Asn^4^-Ala mutant of SLN for this same reason. However, the direct observation of a SLN oligomer required that we image cocrystals of wild-type SLN. Images from frozen-hydrated co-crystals displayed computed diffraction to 15 Å, which improved to approximately 8.5 Å with the averaging of multiple datasets in the p22121 space group (**Table 1** & **Figure 4A**). The projection map for wild-type SLN revealed an oligomeric density (**Figure 4B**) that was similar to previous observations for PLN (9,10). Yet, despite the quality of the data, it was unclear how many SLN molecules were present in the oligomer density. A comparison of the frozen-hydrated projection maps for SERCA-PLN and SERCA-SLN crystals revealed a slightly more compact density for SLN. An interesting difference in the SERCA-SLN crystals was the presence of an additional density adjacent to transmembrane segment M3 of SERCA (**Figure 4C**). The additional density was well resolved in the projection map for wild-type SLN co-crystals, which included 34 images and an excellent overall phase residual (**Table 1**). Moreover, this feature was uniquely observed in wild-type SLN co-crystals and has not been observed in any co-crystals containing PLN, including wild-type and the Ile^40^-Ala and Lys^27^-Ala mutants of PLN (9,10).

**Figure 4:**
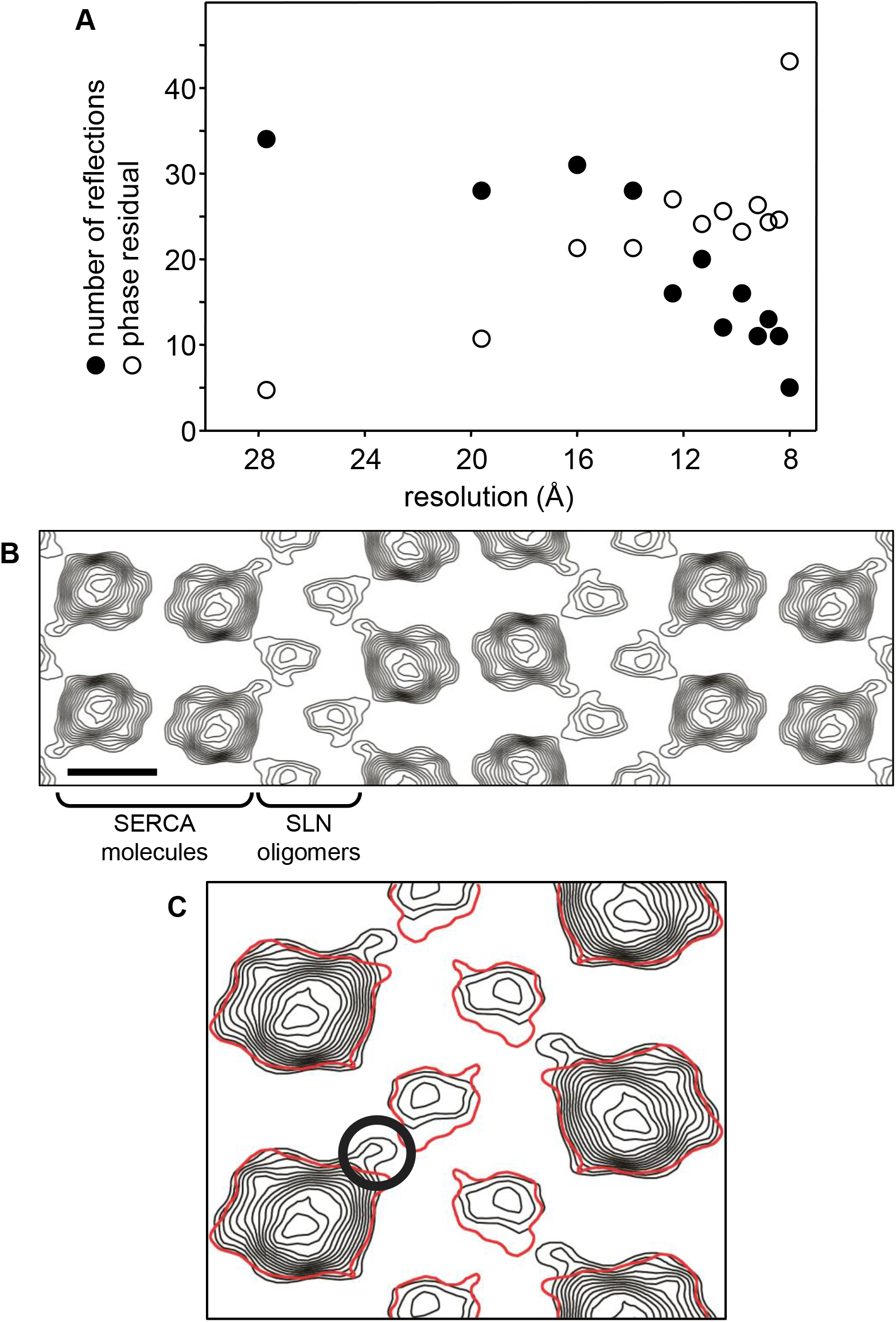
Projections maps of frozen-hydrated two-dimensional crystals of SERCA and SLN. **(A)** Statistics for merging 34 projection images from two-dimensional crystals of SERCA in the presence of wild-type SLN. The number of reflections (*filled circles*) and phase residual in degrees (*open circles*) are plotted versus resolution shell. The data completeness and low phase residuals indicate a resolution of approximately 8.5 Å (random phase residual is 45°, which occurs in the 8Å resolution shell). **(B)** The projection map from 2D crystals of SERCA in the presence of wild-type SLN. The projection map is contoured showing only positive densities (continuous lines; each contour level corresponds to 0.25 σ). For additional crystallographic data, see Table 1. The scale bar corresponds to 50 Å. **(C)** Comparison of the projection maps for 2D crystals of SERCA in the presence of PLN (Glaves et al. 2011) and in the presence of SLN (herein). The same density threshold is shown for SERCA in the presence of PLN (*red contour*) and SERCA in the presence of wild-type SLN (*black contours).* A unique, additional density observed only for SERCA in the presence of SLN is indicated (*circle).* The maximum contour level for the additional density is 0.75 σ, where the SLN pentamer and SERCA had maximum densities of 1.0 σ and 3.25 σ, respectively.

**Table 1:** Summary of crystallographic data for frozen-hydrated two-dimensional co-crystals of SERCA and SLN.

|  | wtSLN |
| --- | --- |
| Number of images | 34 |
| Cell parameters | a = 346.8 Å |
|  | b = 70.5 Å |
| | $\gamma = 90.2^\circ$ |
| Weighted phase residual | 14.8° |

We next sought a more definitive interpretation of the density associated with the SLN oligomer. As a first approach, we integrated and compared the densities corresponding to the PLN pentamers and SLN oligomers in their respective co-crystals. The total density in projection maps corresponding to the PLN pentamer (Figure 6 in (9)) and the SLN oligomer (**Figure 4**) were calculated (**Table 2**). These values were compared to the molecular weights for SLN and PLN. Since the density and molecular weight ratios were in excellent agreement and the PLN density is known to be a pentamer (9), we concluded that the oligomeric state of SLN was consistent with a pentamer. To better visualize the SLN pentamer, we enhanced the high-resolution terms of the cryo-EM density projection map by applying a negative *B*-factor (temperature factor) of 500 Å^2^ during Fourier synthesis (**Figure 5**). This is a standard approach for restoring high resolution information in cryo-EM density maps. The SLN density resolved into a five-lobed pentamer following the truncation of Fourier data at 8.5 Å to minimize the contribution of noise in the map and temperature factor sharpening to improve the contrast and detail for the SLN density. Compared to PLN (9), there were shared and unique features of the SLN projection map. As shared features, both the PLN and SLN densities interacted with the same region of SERCA (transmembrane segment M3) and deviated from the five-fold symmetry expected for a pentamer. As unique features, the SLN density was clearly more pentameric in shape and had a central depression consistent with a central pore. There was an additional density that appeared to connect SERCA and the SLN pentamer, which was consistent with a SLN monomer bridging the interaction.

**Figure 5:**
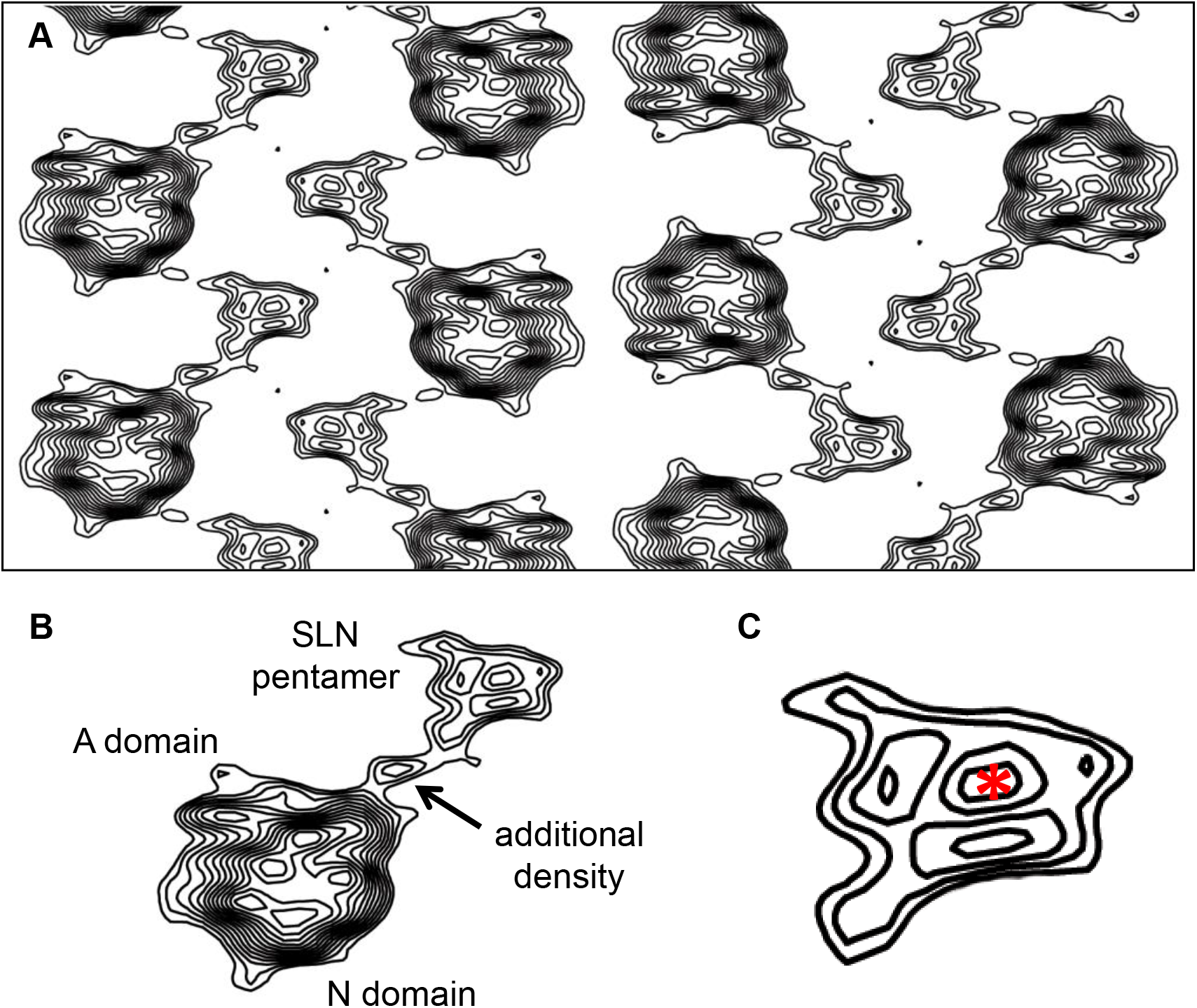
The projection map for SERCA in the presence of wild-type SLN recalculated with an applied negative *B*-factor. **(A)** In the sharpened projection map, the contrast and level of detail is enhanced for both SERCA and SLN. **(B)** A close up view of the density associated with SLN. The size and shape of the SLN densities are now compatible with a pentamer, and there is an additional density adjacent to transmembrane segment 3 (M3) of SERCA. **(C)** The overall shape of the SLN density is consistent with a pentamer, though individual monomers are not resolved. There is a central depression in the SLN density (*asterisk*), further supporting pentameric architecture.

**Table 2:**
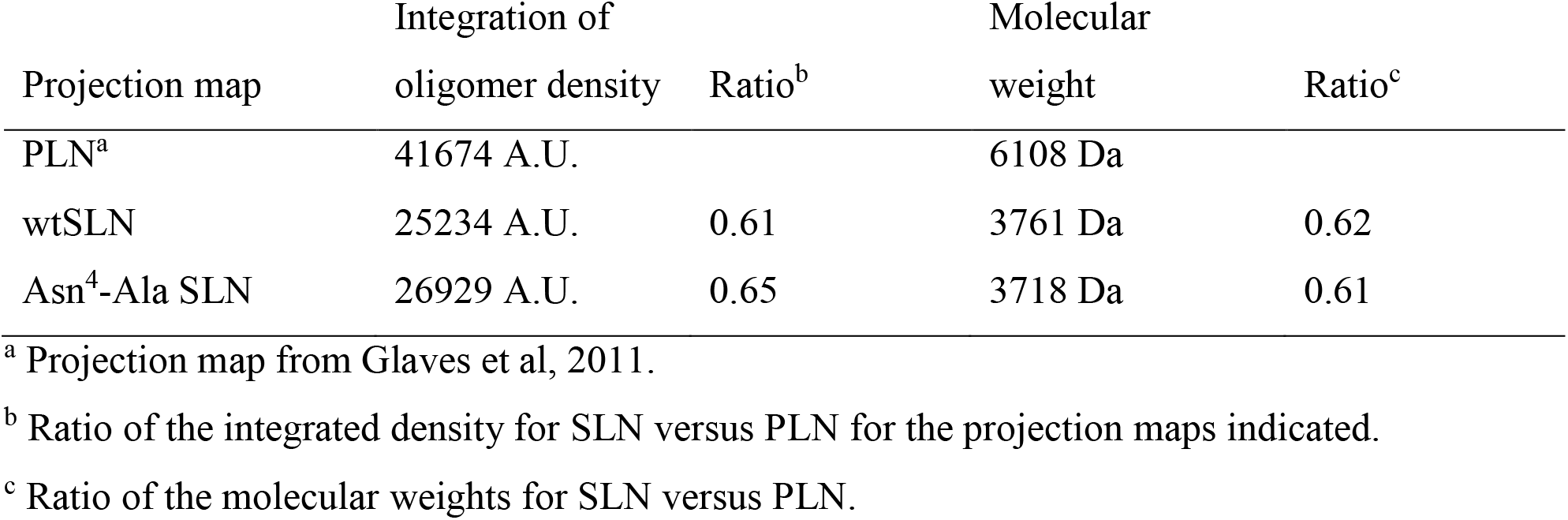
Densities associated with PLN and SLN oligomers.

### Molecular model of the SERCA-SLN complex

Molecular models of the SLN pentamer and the SERCA-SLN heterodimeric complex were generated using protein-protein docking and MD simulations to examine these complexes without the constraints of crystal contacts. For the SLN pentamer, the initial model was constructed based on the symmetric structure of the PLN pentamer (PDB ascension code 2KYV (40)); however, the relaxed structure following MD simulation deviated from the expected structure of a symmetric SLN pentamer (**Figure 6A**). The molecular model was elongated and asymmetric, and appeared to be a complex between a SLN dimer and a SLN trimer. Nonetheless, the *en face* view of the molecular model for the SLN pentamer (**Figure 6B**) closely matched the density for the SLN pentamer in the projection map from 2D crystals (**Figure 5C**). The Leu-Ile repeat that forms the core of the PLN pentameric assembly is relatively conserved between SLN and PLN (**Figure 1**). The SLN residues involved include Leu^10^, Ile^14^, Ile^17^, Leu^21^, Leu^24^, and Ser^28^ and they compare favorably with the PLN resides implicated in pentamer formation (Ile^33^, Leu^37^, Ile^40^, Leu^44^, Ile^47^, and Leu^51^). However, the SLN residues formed complementary interfaces that separately stabilized a SLN dimer and trimer (**Figure 6C**), which together form the SLN pentamer. The model offers an explanation for the reduced stability of the SLN pentamer compared to PLN, and the observation of lower molecular weight species by SDS-PAGE (i.e. monomers & dimers rather than a pentamer). The arrangement of the complementary interfaces in the molecular model also suggests that SLN may be capable of forming an additional oligomeric assembly, such as a hexamer.

**Figure 6:**
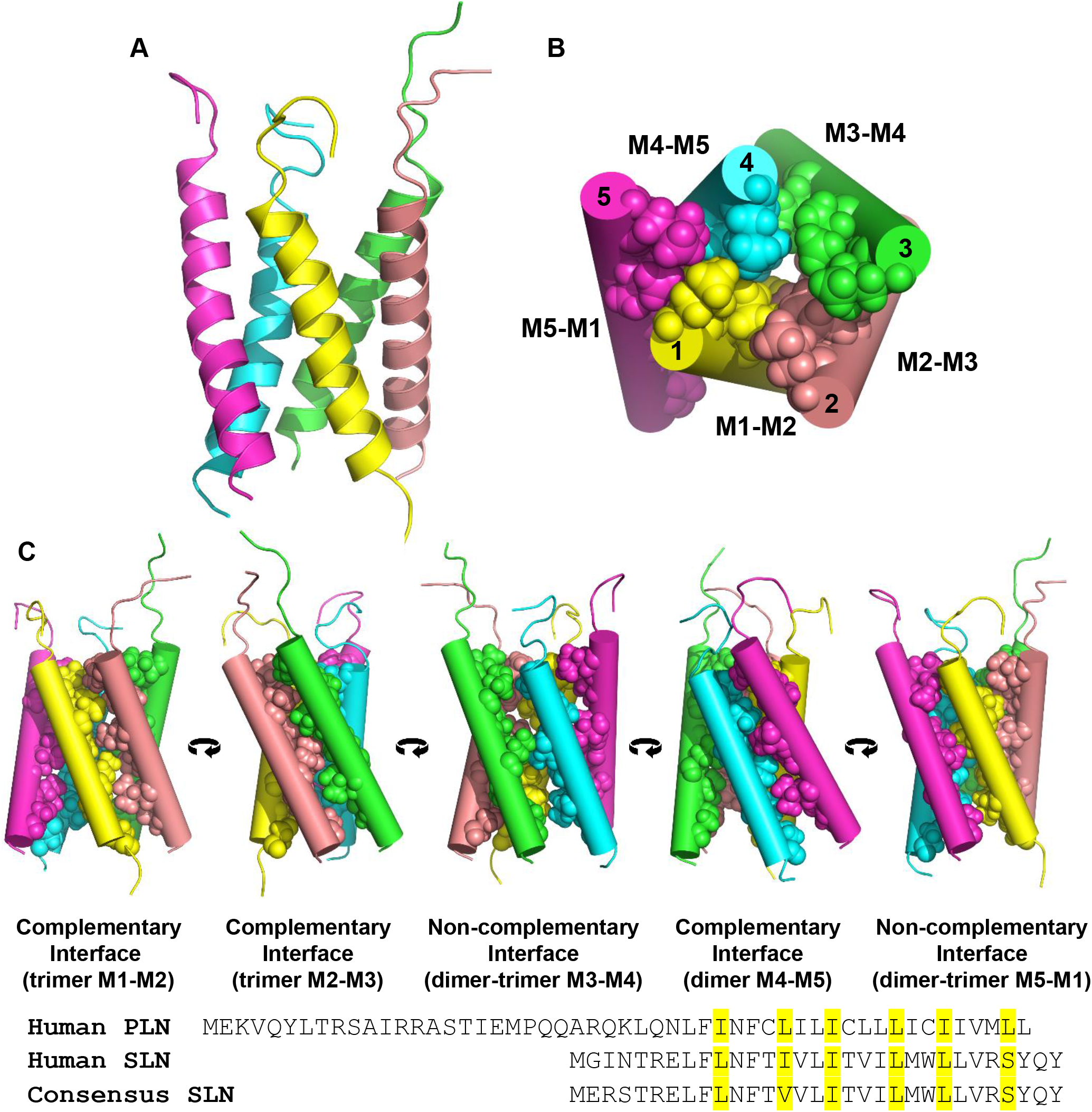
Molecular model for the SLN pentamer. **(A)** Side view of the model with the SLN monomers shown in cartoon format. **(B)** *En face* view of the SLN pentamer with the helices shown as cylinders and residues Leu^10^, Ile^14^, Ile^17^, Leu^21^, Leu^24^, and Ser^28^ shown as spheres. and the SERCA-SLN complex (B) generated from protein-protein docking and MD simulations. M1, M2, and M3 appear to form a trimer, and M4 and M5 appear to form a dimer. **(C)** Side views of the SLN pentamer showing each of the monomer-monomer interfaces. M1-M2, M2-M3, and M4-M5 form complementary hydrophobic interfaces. The non-complementary interfaces between M3-M4 and M5-M1 delineate the SLN dimer and trimer that come together to form the pentamer. A sequence alignment is shown for human PLN, human SLN, and the consensus SLN sequence. The pentamer interface residues are highlighted.

The SERCA-SLN heterodimeric complex was constructed based on the interaction of a SLN monomer with transmembrane segment M3 of SERCA in the 2D crystals (**Figure 7**). Following protein-protein docking, the SERCA-SLN complex that most closely matched the arrangement in the 2D crystals was selected. This complex was embedded in a lipid bilayer, fully hydrated to mimic physiological conditions, and subjected to 400 ns MD simulations. These conditions allowed for the formation of a stable complex with appropriate packing constraints at the SERCA-SLN molecular interface (**Figure 8**). We found that transmembrane segment M3 of SERCA formed a complementary hydrophobic interface with a SLN monomer. The interaction involved the opposite face of SLN’s transmembrane helix (residues Leu^8^, Phe^12^, Leu^16^, & Ile^20^) and the central region of transmembrane segment M3 of SERCA (Leu^266^, Val^269^, & Leu^273^). In addition, Asn^4^ and Arg^6^ of SLN were positioned near a cluster of negatively charged residues on M3 of SERCA (Asp^254^, Glu^255^, & Glu^258^), which is comparable to the interaction between Arg^25^ of PLN and Glu^258^ of SERCA (11). There is limited data on mutagenesis of these residues and the impact on SLN function, with the exceptions being Asn^4^-Ala (**Figure 1**) and Leu^8^-Ala (41). The Asn^4^-Ala mutation results in gain of function, while the Leu^8^-Ala results in loss of function. A lipid straddles the SERCA-SLN interaction on the luminal side of the membrane. Trp^23^ of SLN interacted with the glycerol backbone of the lipid, appearing to position the lipid headgroup toward the M3-M4 loop of SERCA (**Figure 8C, arrow**). This interaction could impact movement of the M3-M4 loop during the calcium transport cycle, thereby decreasing the turnover rate of SERCA.

**Figure 7:**
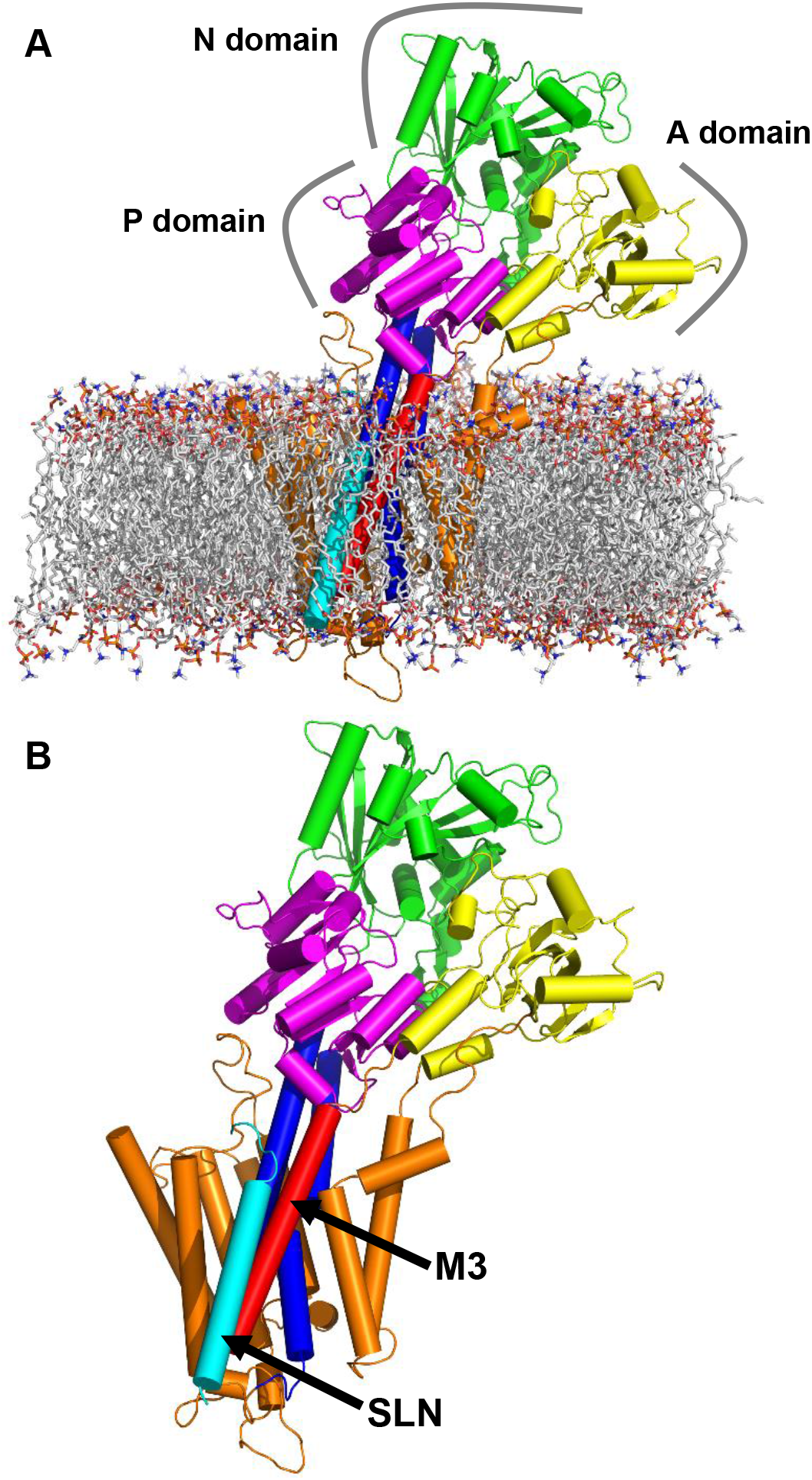
Molecular model for the interaction of SERCA with a SLN monomer. Shown are the side view of the full atomistic model **(A)** and the SERCA-SLN complex **(B)** generated from protein-protein docking and MD simulations. The lipids are shown in stick representation. SERCA and SLN are in cartoon cylinder representation. SLN is in the foreground (*cyan*) adjacent to transmembrane segment M3 of SERCA (*red).* SERCA is colored according to the domain architecture. The nucleotide binding (N domain; *green*), actuator (A domain; *yellow*), and phosphorylation (P domain; *magenta*) domains are indicated. Also indicated are M3 (*red*), M4 and M5 (*blue*), and SLN (*cyan).*

**Figure 8:**
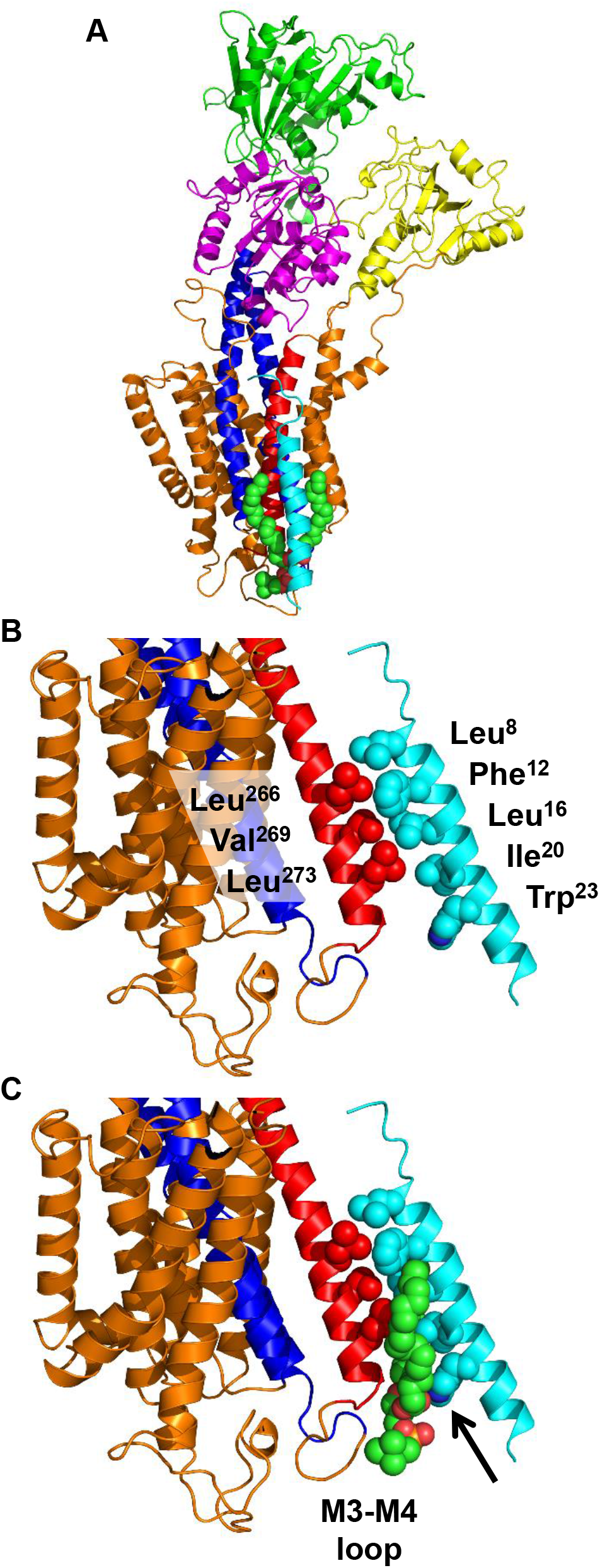
Interaction interface between transmembrane segment M3 of SERCA and a SLN monomer. **(A)** SERCA and SLN are shown in ribbon representation. SERCA is colored according to the domain architecture, with the N domain (*green*), A domain (*yellow*), and P domain (*magenta*) indicated. Also indicated are M3 (*red*), M4 and M5 (*blue*), and SLN (*cyan).* A lipid that straddles the interface is shown in sphere representation (*green).* **(B)** The interface between transmembrane segment M3 of SERCA (*red*) and the SLN monomer (*cyan*) is shown. The residues involved in the interaction include Leu^266^, Val^269^, and Trp^272^ of M3 and Leu^8^, Phe^12^, Leu^16^, and Ile^20^ of SLN. The same interface is shown in **(C)** with the lipid that straddles the interface on the luminal side of the membrane (spheres representation). The acyl chains straddle each side of the SERCA-SLN interface, and Trp^23^ of SLN interacts with the glycerol backbone of the lipid and appears to cause the lipid headgroup to impinge on the M3-M4 loop.

## DISCUSSION

### The PLN pentamer associates with SERCA

The initial model for SERCA inhibition involved reversible binding of monomeric PLN and dynamic equilibrium of the monomer between the SERCA-bound and pentameric states (42). In this scenario, the pentamer was considered an inactive storage form of PLN. However, active roles have been proposed for the PLN pentamer including the modulation of SR cation homeostasis (43) and PKA-mediated phosphorylation (44,45). The PLN pentamer has also been found to stimulate the V_max_ of SERCA (11,15,31,46–48), which depends on the SERCA-PLN molar ratio and density in the membrane (11,46). Providing a context for this latter effect was the finding that PLN pentamers associate with SERCA (9–11). We concluded that the PLN pentamer spontaneously associates with SERCA at a site distinct from the inhibitory groove (M2, M6, & M9 of SERCA (7,49)). The interaction provides an explanation for the stimulation of SERCA’s maximal activity and a molecular model of the complex has been presented (11)

### Functional comparison of SLN and PLN

Does SLN, a PLN homologue, interact with and modulate SERCA maximal activity in a similar manner? To address this question, we focused on large 2D crystals of the SERCA-SLN complex. We characterized wild-type SLN and a gain-of-function mutant deemed suitable for structure determination. Prior mutagenesis studies of SLN targeted select residues homologous to either the functional interface or the pentamer interface of PLN (33). Essential functional residues such as Leu^8^ and Asn^11^ (Leu^31^ and Asn^34^ of PLN) were found to cause loss of function when mutated to alanine, yet mutagenesis revealed little similarity to the pentamer interface of PLN. Nonetheless, SLN has since been shown to form a pentamer (12,13). In the cytoplasmic domain of SLN, only Thr^5^ was studied as a potential site of regulation by phosphorylation. Sequence comparison with PLN indicated that the cytoplasmic domain of SLN consists of the first seven residues (**Figure 1**; ^1^MGINTRE^7^ in humans; ^1^MERSTQE^7^ consensus sequence), followed by the transmembrane domain (residues 8-26), and a luminal extension (residues 27-31).

In the present study, we mutated Asn^4^ of human SLN to alanine (Asn^4^-Ala) and found it to result in gain of function. Asn^4^ of SLN is similar to Lys^27^ of human PLN, and both residues deviate from the consensus sequence and are unique to humans (and primates). The gain-of-function behaviour observed for both residues suggests commonality in function between the membrane proximal region of PLN and the N-terminus of SLN, despite low overall sequence conservation. The crystal structure of the SERCA-PLN complex (49) places this residue (Asn^27^ in canine PLN; Lys^27^ in human PLN) between Phe^809^ and Trp^932^ of SERCA, which draws this region of PLN closer to M6 and M9 of SERCA. The homologous consensus residue in the crystal structures of the SERCA-SLN complex (7,8) is Ser^4^ of rabbit SLN. This residue does not interact with Phe^809^ and Trp^932^ of SERCA, which allows a small rotation of the SLN helix in this region. However, this residue is Asn^4^ in human SLN and there is functional similarity to Asn^27^ in canine PLN and Lys^27^ in human PLN. Thus, Asn^4^ in human SLN may interact with Phe^809^ and Trp^932^ of SERCA, thereby playing a role in positioning the N-terminus of SLN. Moreover, Asn^4^ in SLN and Lys^27^ in PLN appear to be evolutionary sequence modifications unique to primates (50).

### SLN as a pentamer

The original mutagenesis studies of SLN did not reveal the expected gain-of-function pattern associated with PLN pentamer destabilization, supporting observations by SDS-PAGE that SLN did not appear to form an oligomeric structure (33). However, several studies have since shown that SLN can form a mixture of oligomeric species including a pentamer (12,13) and that mutations can cause SLN depolymerisation (13). Intriguingly, recent studies have reported chloride and phosphate transport properties for SLN (51,52) and potassium transport properties for PLN (53), which would be consistent with a common oligomeric architecture for the two proteins. Here, we report the first direct observation of a SLN pentamer in 2D crystals with SERCA. Both the PLN pentamer characterized previously (9) and the SLN density observed in this study (**Figure 5**) deviate from the five-fold symmetry expected for a pentamer, though the SLN density is more symmetric. The SLN density also displayed a distinct central depression (**Figure 5C**) consistent with a channel-like architecture, which was not as well defined for the PLN pentamer (9). The distortion of symmetry for both the PLN and SLN pentamers may be due to noise and the moderate resolution (8.5 Å) of the projection maps, or it may be a modulation of the PLN and SLN pentamers due to the physical interaction with SERCA. This latter point seems plausible, given that molecular dynamics simulations of the SERCA-PLN complex revealed that the symmetry of the PLN pentamer is altered in the interaction with SERCA (11).

A molecular model of the SLN pentamer was generated by protein-protein docking and MD simulations. An initial symmetric model of the SLN pentamer was generated based on docking of SLN monomers to the PLN pentamer (40). Following MD simulations, the final model of the SLN pentamer was asymmetric and elongated (**Figure 6B**), which agreed well with the density map from 2D crystals (**Figure 5C**). Moreover, the SLN pentamer appeared to be a complex of a SLN dimer and a SLN trimer, with complementary hydrophobic interfaces that stabilize the dimer and trimer (**Figure 6C**).

### Interaction between SERCA and SLN

As with PLN, an interaction between an oligomeric form of SLN and transmembrane segment M3 of SERCA was observed in the 2D crystals. This interaction site does not correspond to the inhibitory groove of SERCA (M2, M6, & M9). This interaction only occurs at a high membrane density and molar excess of PLN relative to SERCA, similar to what is found in cardiac SR membranes (36,48,54–57). This packing density of SERCA and PLN is also similar to what is found in large 2D crystals (9,10), albeit without the regular order induced by the crystal lattice. The molar ratio of SERCA-SLN in the reconstituted proteoliposomes, 1:5, is higher than that found in skeletal muscle SR membranes (33). At these ratios and membrane densities, SLN and PLN monomers and oligomers will be in close proximity to SERCA molecules in their respective SR membranes. Given their close proximity, it is not surprising that the oligomeric forms of these proteins are capable of a physical association with SERCA. In the case of PLN, we found that the interaction of the PLN pentamer with M3 of SERCA correlates with an increase in the V_max_ of SERCA (11). The proximity of this interaction to the calcium entry funnel of SERCA provided an explanation for the increased turnover rate of SERCA – namely, the cytoplasmic domains of PLN lie along the membrane surface and perturb lipid packing adjacent to the calcium access funnel.

The interaction of a pentamer with the M3 accessory site was also observed for 2D crystals of SLN and SERCA. The PLN and SLN pentamers occupy a similar position, though the SLN pentamer has a narrower density profile reflecting its smaller size (**Figure 4C**). While the PLN pentamer directly contacts M3 of SERCA, the SLN pentamer is more distant from M3 and an additional density bridges the interaction (**Figure 5B**). The additional density has the appearance of an α-helix oriented perpendicular to the membrane plane as visualized by cryo-electron crystallography (58). This is consistent with a SLN monomer lying adjacent to M3 of SERCA. The question then becomes, why do PLN and SLN interact with this region of SERCA? In the model of the SERCA-PLN pentamer complex, the interaction appears to be stabilized by electrostatic interactions between the cytoplasmic domain of PLN and transmembrane segment M3 of SERCA. Indeed, there is a cluster of negatively charged residues on M3 of SERCA at the cytoplasmic membrane surface (Asp^254^, Glu^255^, & Glu^258^). The cytoplasmic domain of PLN is highly basic and a stable electrostatic interaction forms between Arg^25^ of PLN and Glu^258^ of SERCA. There are additional negatively charged residues on M1 along the calcium access funnel of SERCA (Glu^51^, Glu^55^, Glu^58^, & Asp^59^). Complementary to this, there are several positively charged residues in the cytoplasmic domain of PLN (Arg^9^, Arg^13^, Arg^14^, & Lys^27^). Thus, the interaction between the PLN pentamer and SERCA may be driven by electrostatic attraction. By comparison, Asn^4^ and Arg^6^ of human SLN are proximal to Glu^258^ of SERCA in a similar manner as Arg^25^ of PLN (the consensus SLN sequence includes Ser^4^, which is Asn^4^ in humans and primates (50)). Thus, SLN may be electrostatically drawn to M3 of SERCA, though this is likely reduced compared to PLN.

In the model of the SERCA-SLN complex, the monomer forms a complementary hydrophobic interface with a central region of transmembrane segment M3 of SERCA (**Figure 8**). This interface, combined with the reduced stability of the SLN pentamer, may explain why a SLN monomer was found associated with the M3 site. As stated above, the residues involved are on the opposite face of the SLN transmembrane helix (Leu^8^, Phe^12^, Leu^16^, & Ile^20^) compared to the residues that stabilize the model of the SLN pentamer (Leu^10^, Ile^14^, Ile^17^, Leu^21^, Leu^24^, & Ser^28^), and the SLN monomer appears to form a more direct interface with M3 of SERCA than the PLN pentamer. The different interactions between PLN and SLN and the M3 site of SERCA correlate with distinct functional effects on the V_max_ of SERCA. The PLN pentamer increases the V_max_ of SERCA, while SLN decreases the V_max_ of SERCA. In the model of the SERCA-PLN pentamer complex, a cluster of negatively charged residues on M3 of SERCA attracts the cluster of positively charged residues in the cytoplasmic domain of PLN. This draws the PLN pentamer toward M3, with an interaction interface that involves two PLN monomers and a lipid acyl chain (11). This places the cytoplasmic domains of PLN proximal to the calcium access funnel of SERCA, without impeding the movement of M3 during the calcium transport cycle. The V_max_ stimulation by the PLN pentamer was attributed to membrane destabilization by the cytoplasmic domain of PLN near the calcium access funnel of SERCA, thereby allowing a more rapid turnover rate (11). This is absent in the SERCA-SLN complex, suggesting that the M3 interaction alone does not stimulate SERCA. Indeed, the model of the SERCA-SLN monomer complex involves a complementary hydrophobic interface, which would be expected to reduce the turnover rate of SERCA by impeding the movement of M3 during the calcium transport cycle. A lipid straddles the SERCA-SLN interface on the luminal side of the membrane and may play a role in mediating the functional effect of SLN. We have previously shown that the unique luminal tail at the C-terminus of SLN (^27^RSYQY^31^ sequence) contributes to the decrease in the V_max_ of SERCA (5). The crystal structures of the SERCA-SLN inhibitory complex (7,8) and the model of the SERCA-SLN complex presented herein (**Figure 7 & 8**), suggest that the luminal tail serves to position the transmembrane helix of SLN in the membrane for optimal interaction with SERCA.

Finally, the full complex that interacts with M3 of SERCA involves a SLN monomer that mediates the interaction between the SLN pentamer and M3 of SERCA, though the role of the SLN pentamer in this complex remains unknown. However, compared to PLN, the lower stability of the SLN pentamer combined with the M3 interaction may select for a SLN monomer. Indeed, the molecular models for the SLN pentamer and the SERCA-SLN interaction raise the possibility that the full complex represents a transition state in which the interaction of M3 with an oligomeric form of SLN causes a SLN monomer to dissociate for optimal interaction with M3. Since both the SERCA-PLN (11) and SERCA-SLN (**Figure 8**) complexes appear to involve a lipid molecule at the interface, this may explain the dependence of the interaction on high membrane densities and low lipid-to-protein ratios.

## AUTHOR CONTRIBUTIONS

JPG, JOP, and PAG performed the research, analyzed data, and contributed to the writing and editing of the manuscript; MJL contributed to the design and analysis of the data, and editing of the manuscript; LMEF performed and analyzed the protein-protein docking and molecular dynamics simulations; HSY designed the research, analyzed the data, and wrote the manuscript.

## ACKNOWLEDGEMENTS

This work was supported by a grant from the Heart and Stroke Foundation of Canada and the National Institutes of Health (R01HL092321) to HSY. JPG and PAG were supported by Canada Graduate Scholarships from the Canadian Institutes of Health Research and Alberta Innovates – Technology Futures. LMEF was supported by a grant from the National Institutes of Health (R01GM120142). MJL was supported by grants from the Canadian Institutes of Health Research and the Natural Sciences and Engineering Research Council of Canada.

---

**Supplemental Figure 1:**
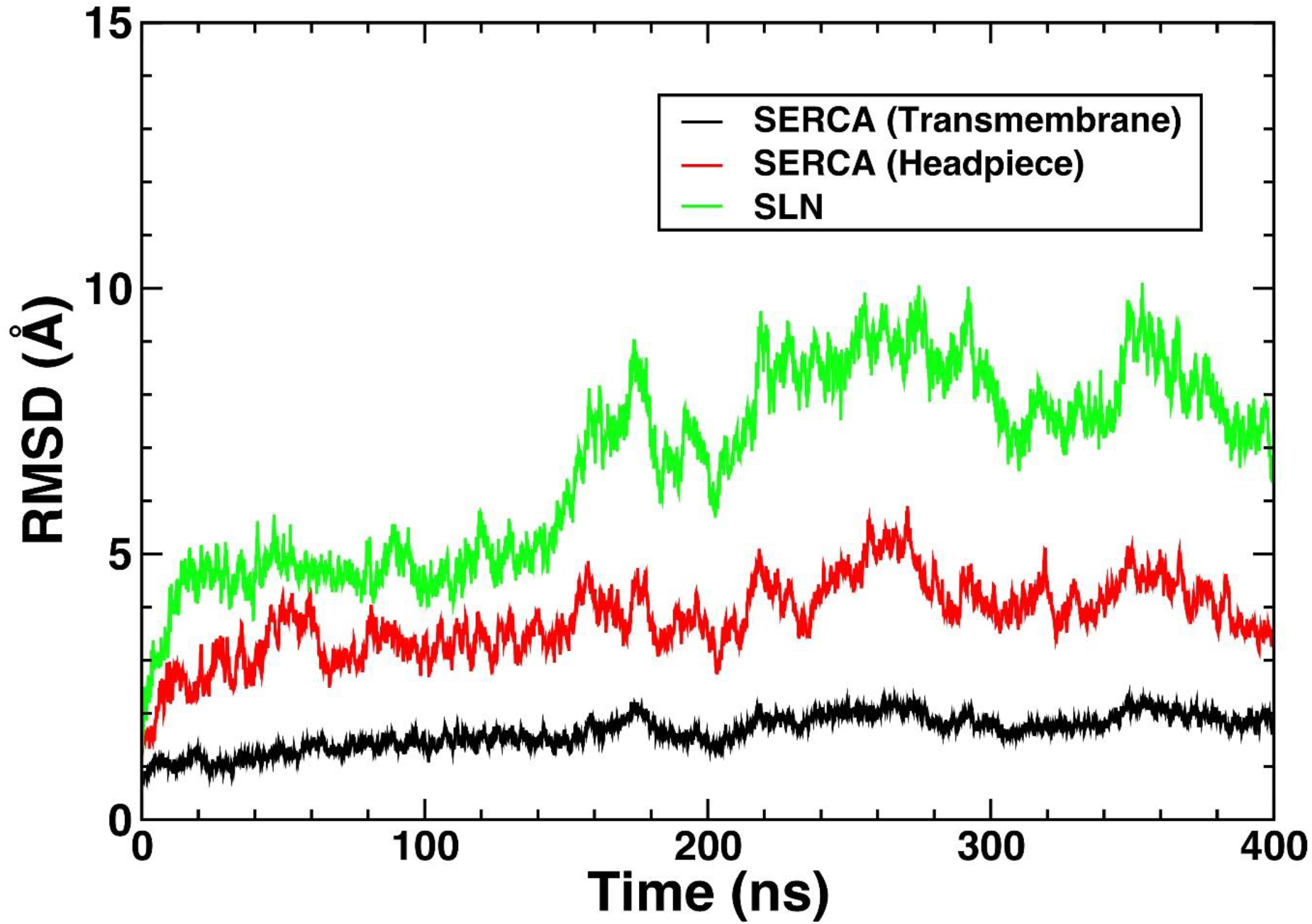
Time-dependent changes in the RMSD for the SLN monomer and SERCA in MD simulations of the heterodimeric complex.

